# Right-sided brain lesions predominate among patients with lesional mania: evidence from a systematic review and pooled lesion analysis

**DOI:** 10.1101/433292

**Authors:** J. Bernardo Barahona-Corrêa, Gonçalo Cotovio, Rui M. Costa, Ricardo Ribeiro, Ana Velosa, Vera Cruz e Silva, Christoph Sperber, Hans-Otto Karnath, Suhan Senova, Albino J. Oliveira-Maia

## Abstract

**Background:** Despite claims that lesional mania is associated with right-hemisphere lesions, supporting evidence is scarce, and association with specific brain areas has not been demonstrated.

**Aims:** To test whether focal brain lesions in lesional mania are more often right-than left-sided, and if lesions converge on areas relevant to mood regulation.

**Methods:** We performed a systematic literature search (PROSPERO registration CRD42016053675) on PubMed and Web-Of-Science, using terms that reflected diagnoses and structures of interest, and lesional mechanisms. Two researchers reviewed the articles separately according to PRISMA Guidelines, to select reports of adult-onset hypomania, mania or mixed state following a focal brain lesion. When available, eligible lesion images were manually traced onto the corresponding slices of MNI space, and lesion topography analyzed using standard brain atlases. Pooled-analyses of individual patient data were performed.

**Results:** Data from 207 lesional mania patients was extracted from 110 reports. Among patients with focal lesions (N=197) more patients had lesions involving the right (84.3%) than the left (34.5%) hemisphere. Among 54 lesion images that were available, right-sided predominance of lesions was confirmed, and found to be was conserved across multiple brain regions, including the temporal lobe, fusiform gyrus and thalamus. These, in addition to several frontal lobe areas, were also identified as preferential lesion sites in comparisons with control lesions.

**Conclusions:** Pooled-analyses, based on the most comprehensive dataset of lesional mania available to date, confirm a preferential association with right-hemisphere lesions, while suggesting that several brain areas/circuits, relevant to mood regulation, are most frequently affected.

## Introduction

Bipolar disorder (BPD), affecting 3-6% of the population worldwide, manifests as a recurrent, episodic disturbance of mood, sleep, behavior, and perception, including at least one episode of acute mania or mixed affective state. While the first manifestations of BPD could be depressive episodes, diagnosis is typically established after the first manic, hypomanic or mixed episode. The overwhelming majority of such episodes are idiopathic, leading to a diagnosis of primary BPD. Secondary mania, in contrast, refers to cases where a manic, hypomanic or mixed episode first appears after an organic insult, including structural brain lesions. Typically occurring at a later age(1), the diagnosis of lesional mania requires temporal proximity between onset of the inaugural manic episode and prior occurrence of an identifiable brain insult. Common causes include stroke, traumatic brain injury, or tumors(1). Although lesional mania has traditionally been associated with right-hemisphere brain lesions(2), the evidence supporting this claim is mostly anecdotal (3). Moreover, while a recent narrative review found that the thalamus, hypothalamus, basal ganglia and frontal and temporal cortices were the most frequent lesion locations(4), it remains unresolved if lesional mania predominantly involves a specific brain area or network.

Studying lesional mania is a valuable approach to understand the neuroanatomy of primary mania and BPD. In fact, the direction of causal associations between brain structure and behavioral changes is clearer for lesional mania than for primary BPD. Furthermore, this approach may highlight brain areas and networks missed by comparative image protocols, the latter being inevitably contaminated by unspecific, non-causal positive findings(5). Here we present the results of a systematic literature review on lesional mania, with pooled analyses of anatomical data reported for individual cases, as well as comparisons of this data with that of lesion distribution in control populations. While the main goal of this pooled analysis was to confirm whether brain lesions in lesional mania are more often right-than left-sided, we further explored whether lesions converged on specific areas or circuits relevant to mood regulation.

## Methods

### Protocol and registration

The protocol was published in PROSPERO database (CRD42016053675) and can be consulted for full method description (http://www.crd.york.ac.uk/PROSPERO/display_record.asp?ID=CRD42016053675).

### Information sources and search strategy

Search was performed on PubMed and Web-of-Science between May 2015 and October 2016. Search terms reflected diagnoses of interest (bipolar disorder, manic, mania), structures of interest (cerebral, cerebellum, brain, central nervous system) and possible mechanisms of lesion (injury, tumor, neoplasm, mass, infection, abscess, cyst, stroke, hemorrhage, bleeding). Filters were applied to restrict search results to adult human subjects (Table S1).

### Study selection and eligibility criteria

After eliminating duplicates, two researchers reviewed the list of articles separately, selecting eligible reports according to PRISMA procedures. Articles in English, French, German, Portuguese or Spanish were considered, regardless of publication date or country of origin. Eligible cases were 18 years or older, with a distinct episode of behavioral change lasting at least four days, and causing significant psychosocial impairment, manifesting with: A – elevated, expansive, or irritable mood and abnormally and persistently increased goal-directed activity or energy; B – at least three of the following: inflated self-esteem or grandiosity, decreased need for sleep, excessive talkativeness, flight of ideas, distractibility, increased goal-directed activity, and excessive involvement in activities with potentially painful consequences(6). Reports that did not provide details on behavioral changes remained eligible if authors explicitly stated that they met contemporary DSM or ICD criteria for manic, hypomanic or mixed affective state. Eligibility further required at least one confirmed brain lesion that preceded the first manic/hypomanic manifestations. Cases where a brain lesion was diagnosed after the first manic/hypomanic manifestations were considered if the lesion was unequivocally acquired prior to BPD onset. Cases were excluded if no brain lesion was identified, the chronology between lesion occurrence and manic symptoms could not be unequivocally established, or the brain lesion occurred after the first manic syndrome. Literature reviews or meta-analyses were excluded, but were screened for additional references, as were reference lists of eligible articles.

### Data extraction, data items and risk of bias

Two researchers extracted data separately according to PRISMA guidelines. For each paper, author name, title and journal, publication year, study type, and number of reported and eligible cases was recorded. For each eligible case we noted age at first episode of lesional mania, gender, hand dominance, time-interval between brain lesion and mania onset, availability of lesion image (MRI, CT, SPECT, drawing on a standard brain atlas, or photographs of autopsy specimens), lesion location and nature as described by original authors, previous history of depression, personal or family history of other neuropsychiatric disorders, and mania symptoms mentioned in the case description. For systematic assessment of study quality we created a Clinical Quality Assessment scale (CQA) and a Brain Lesion Documentation Assessment scale (BLDA, see Table S2). Eligible lesion images, i.e. BLDA≥3, were manually transcribed onto the corresponding slices of the MNI_ICBM152NLin2009 atlas (http://www.bic.mni.mcgill.ca/ServicesAtlases/ICBM152NLin2009), using MITK software v2014.10.00 (http://mitk.org/wiki/MITK). Two of the authors, including a neuroradiologist, performed this task jointly, and a third author, who is a neurosurgeon, then independently reviewed lesion traces. When tracing, each of five group diagrams comprising a total of 54 subjects were considered as single cases.

### Summary measures and synthesis of the results

Based on descriptions by the original authors and/or available brain images, we classified each lesion according to laterality and brain region(s). For traced lesion images, quantification of lesion distribution in grey-matter (GM) and white-matter (WM) was performed on Anatomist software (http://brainvisa.info/web/download.html), using the Automated Anatomical Labeling atlas (AAL; http://www.cyceron.fr/index.php/en/plateforme-en/freeware), and the John Hopkins University (JHU) WM tractography atlas (http://cmrm.med.jhmi.edu/), respectively.

### Statistical analysis

Data are presented as % patients or mean ± standard deviation (SD). To test the null hypothesis that lesions were randomly distributed across both brain hemispheres, and because patients could have bilateral lesions, we used McNemar’s test for repeated measures to compare the proportion of patients with left- vs. right-sided lesions in the entire sample with focal lesions, first for the whole brain and then for each pre-defined region of interest. In traced lesion images, we further compared, for each area of the AAL and JHU atlases, the mean proportion of affected voxels on the left- vs. right-hemisphere, using Wilcoxon’s signed-rank test. Furthermore, we compared lesion distribution among our tumor sub-sample with data for 169934 adult-onset brain tumours reported in a database published by Ostrom et al(7). To do so we classified tumors in our database according to the criteria used by Ostrom et al, and compared the proportions of patients with lesions in each brain region with those reported by Ostrom et al, using Fisher’s exact tests. Similarly, for cases of right-sided vascular lesions with traced lesion images, we compared the proportion of patients with lesions in each area of the GM and WM atlases to that in a sample of 439 unselected right-hemisphere stroke patients(8). Here, the presence of a lesion in a particular brain area was defined according to a thresholds relating to the proportion of the total volume of that area that was lesioned. For lesional mania cases, where only few brain-image slices were available in each patient, a lesion was defined when more than 0.1%, and at least 5 voxels, were lesioned in a particular area. In controls, where we had access to full MRI scans, the threshold was defined as 15% of lesioned voxels. According to these thresholds, both groups had a similar mean number of affected brain areas per individual (8.99 ± 8.8 and 9.5 ± 7.4 respectively, p=0.8). For all analyses, statistical significance was defined according to Benjamini-Hochberg(9), assuming a false discovery rate (FDR) of 0.1.

## Results

### Literature review

Literature review identified 110 eligible articles (Fig. 1), comprising 207 case-descriptions, including both focal – involving at least one circumscribed brain area(10) – and diffuse lesions (where damage is spread over wide or multiple brain areas). Brain lesion documentation was provided for 115 patients (55.6%), namely from MRI (n=28), CT (n=29), schemes/drawings (n=55), and autopsy photographs (n=3). Forty-nine of these cases were traced on the MNI atlas. Seven were not eligible due to low image quality (BLDA≤2) and lesions were had diffuse. Lesion tracings obtained from 5 diagrams depicting several lesions jointly, and representing a total of 54 patients, were further considered for lesion topography analyses.

**Figure 1 –.**
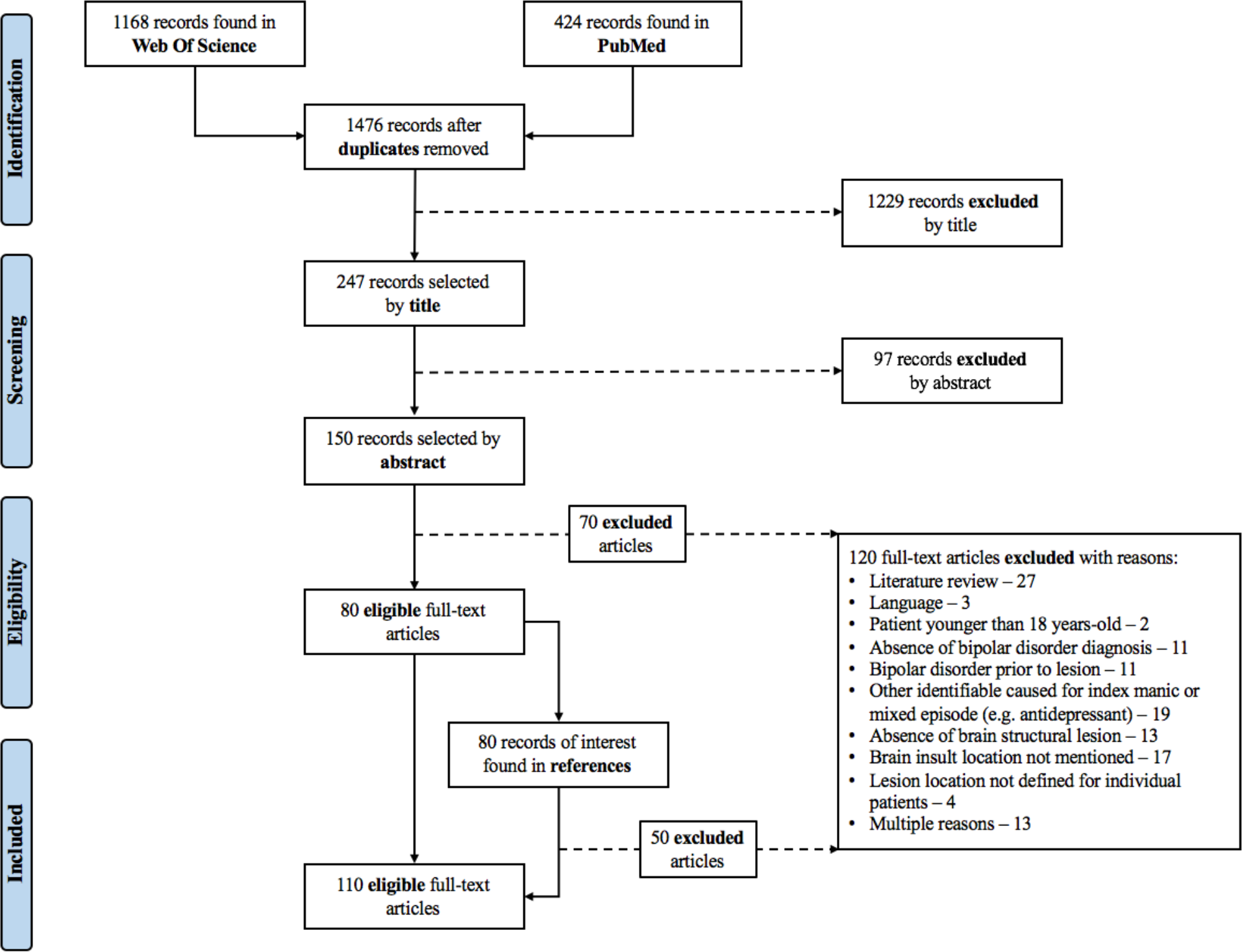
Article selection flowchart (according to PRISMA Statement)

### Results and synthesis of studies

Mean age at lesional mania onset was later than what is typically reported for primary BPD [48.6±17.5 vs. 20.2±11.8 years in Morken et al(11), for example], and most patients were male (63.7%) and right-handed (87.6%). A prior history of depression was mentioned in 14.2% of patients, and 27.6% had another neuropsychiatric diagnosis, epilepsy being most common. Just over half of the cases developed manic symptoms in the first month after the respective brain lesion. Most cases were vascular lesions (51.3%) or tumors (20.4%). Comparison between patients with vascular and non-vascular lesions revealed that the former were older, more frequently right-handed and had a less frequent history of depression (table 1).

**Table 1–.** Demographic and clinical data extracted from eligible cases

| Characteristic |  | Total Sample<br>(N=207) | Vascular vs. Non-Vascular Etiology<sup>a</sup> |  |  |
| --- | --- | --- | --- | --- | --- |
|  |  |  | Vascular (N=74) | Non-Vascular<br>(N=69) | p value |
|  |  | Mean ± SD (N)<br>or N (%) | Mean ± SD (N)<br>or N (%) | Mean ± SD (N)<br>or N (%) |  |
| **Age at onset** |  | 48,6 (+/-17,5; 191) | 56,2 (+/-16,7; 54) | 43,4 (+/-15,1; 68) | <0,0001 |
| **Gender** | Female | 73 (36,3) | 22 (32,4) | 25 (36,2) | n.s. |
|  | Male | 128 (63,7) | 46 (67,7) | 44 (63,7) |  |
| **Hand Dominance** | Right | 92 (87,6) | 31 (96,9) | 9 (69,2) | 0,02 |
|  | Left/Amb. | 13 (12,4) | 1 (3,1) | 4 (30,8) |  |
| **Time E-MM** | ≤1m | 37 (53,6) | 24 (63,2) | 13 (41,9) | n.s. |
|  | >1m | 32 (46,4) | 14 (36,8) | 18 (58,1) |  |
| **Aetiology** | Vascular | 98 (51,3) |  |  |  |
|  | Tumour | 39 (20,4) |  |  |  |
|  | TBI | 24 (12,6) |  |  |  |
|  | Other | 30 (15,7) |  |  |  |
| **Previous Depression** | Yes | 23 (14,2) | 7 (11,1) | 11 (20,8) | n.s. |
|  | No | 139 (85,8) | 56 (88,9) | 42 (79,25) |  |
| **Personal history of other NP disorder** | Yes | 45 (27,6) | 8 (13,8) | 21 (35,6) | 0,01 |
|  | No | 118 (72,4) | 50 (86,2) | 38 (64,4) |  |

<sup>a</sup> Does not include the following case series, which did not provide enough information on individual lesion
Amb. – Ambidextrous; E – Event causing brain insult; m – month; NP – neuropsychiatric.

### Anatomical distribution of lesions

Among 197 cases with focal brain lesions, these were exclusively right-sided in 61.4%, exclusively left-sided in 11.7%, bilateral in 22.8% and midline in 4.1% (p<0.0001, McNemar’s test). Thus, 166 patients (84.3%) had right-hemisphere lesions, while only 68 (34.5%) had left-sided lesions. Further comparisons demonstrated a significantly higher proportion of lesions on the right, relative to left, frontal (p=0.0001), temporal (p=0.004), parietal (p=0.01), and occipital lobes (p=0.02), as well as thalamus (p=0.0001), basal ganglia (p=0.001) and subcortical WM (p=0.03; McNemar’s tests; FDR corrected; Fig. 2A).

**Figure 2 –.**
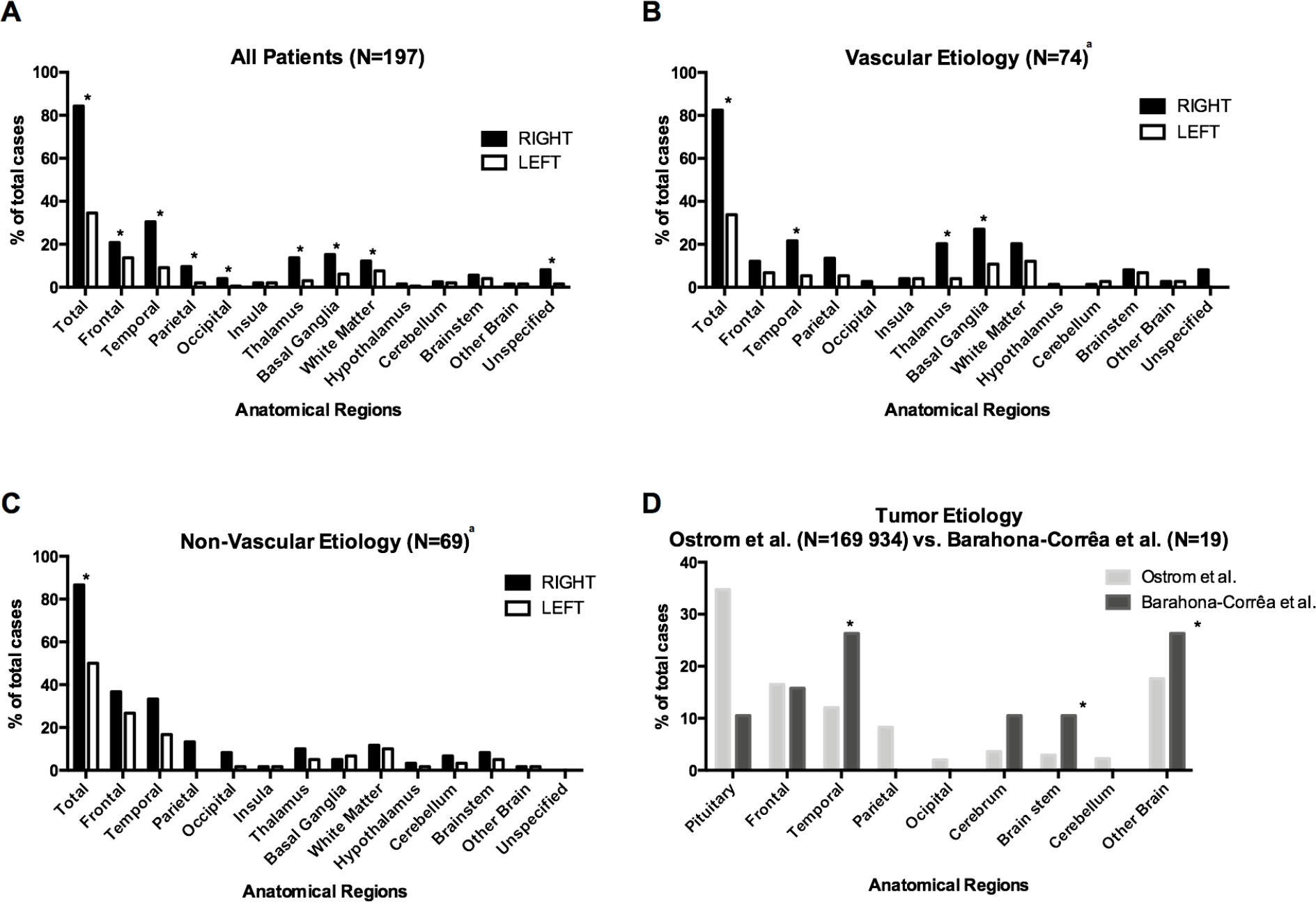
Lesion distribution by major brain areas. A) Lesion distribution for all cases in the literature review. (B) Lesion distribution for cases with vascular lesions. (C) Lesion distribution for cases with non-vascular lesions. (D) Comparison of lesion distribution for tumor cases identified in this literature review (n=19) with tumor distribution described by Ostrom et al for a large patient database (n=169934)(7). In what brain tumors are concerned, regions were defined according to the International Classification of Diseases for Oncology (ICD-O), without considering tumours originating from the meninges (n=14), ventricles (n=2), cranial nerves (n=1), or of unspecified origin (n=4). “Other Brain” refers to lesions spanning multiple brain areas (C71.8: “*neoplasm involving two or more sites*, *corpus callosum and tapetum”*) or when areas were insufficiently specified (C71.9: intracranial site, cranial fossa not otherwise specified, anterior cranial fossa, middle cranial fossa, posterior cranial fossa and suprasellar). “Cerebrum” refers to basal ganglia, central white matter, unspecified cerebral cortex, cerebral hemisphere, cerebral white matter, corpus striatum, globus pallidus, hypothalamus, insula, internal capsule, island of Reil, operculum, pallium, putamen, rhinencephalon, supratentorial brain not otherwise specified and thalamus (C71.0). ^a^ Does not include the following case series, which did not provide enough information on individual lesion etiology: Carran 2003, Robinson 1988, Starkstein 1987 and Starkstein 1991 (See Supplementary Material for complete references – Table S3). *p-value<0.05

Separate comparisons for vascular and non-vascular lesions confirmed an overall predominance of right-sided lesions, in both cases, and for the temporal cortex, basal ganglia and thalamus for vascular lesions only (Fig. 2B and 2C). In quantitative GM and WM analyses of traced lesions (Fig. 3), when compared to the corresponding areas in the left hemisphere, a significantly higher mean proportion of lesioned voxels was found for the right hippocampus, parahippocampal, middle temporal, inferior temporal, lingual and fusiform gyri, caudate, putamen, thalamus, internal capsule, posterior thalamic radiation, sagittal stratum and fornix (Tables 2&S4).

**Figure 3 –.**
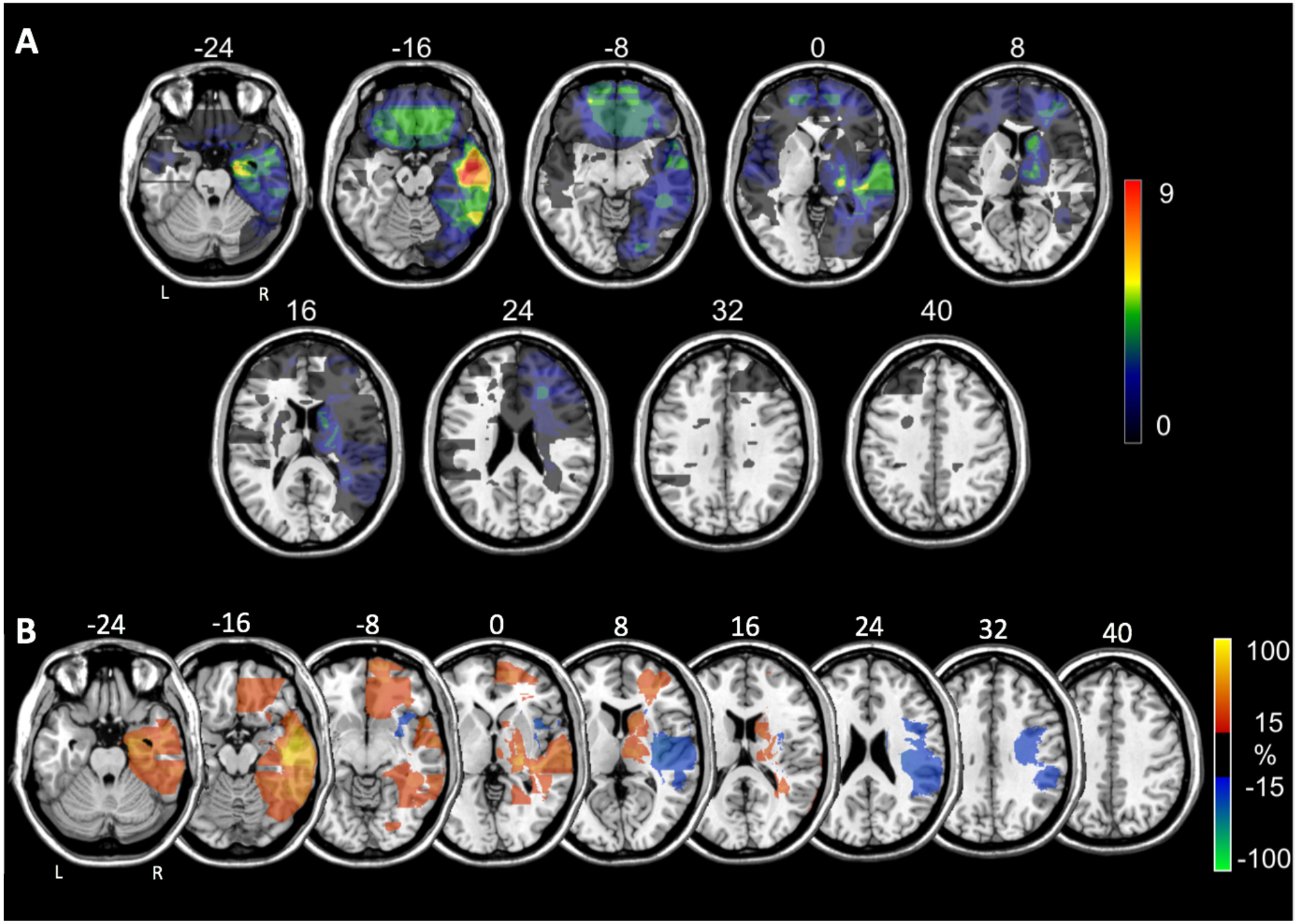
Distribution of brain lesions associated with secondary bipolar disorder in 54 patients with eligible lesion images. (A) Comparison between right vs. left-sided lesions. Each lesion was traced manually onto a common brain atlas (MNI) and projected on the closest depicted slice. Numbers above slices indicate z-coordinates in MNI space. Number above the color bar indicates maximal number of lesions overlapping on a single voxel. (B) Subtraction plot contrasting 27 right-sided stroke patients with secondary bipolar disorder (red-yellow) versus 439 unselected right hemisphere stroke patients (blue-green). In this plot, a value of, for example, 30, reflects that the voxel is damaged 30% more frequently in bipolar patients than in unselected patients (for more details on the method see Rorden & Karnath, 2004(5)). To improve visualization, lesions of bipolar patients were projected onto the closest depicted slice before plot generation.

**Table 2 –.**
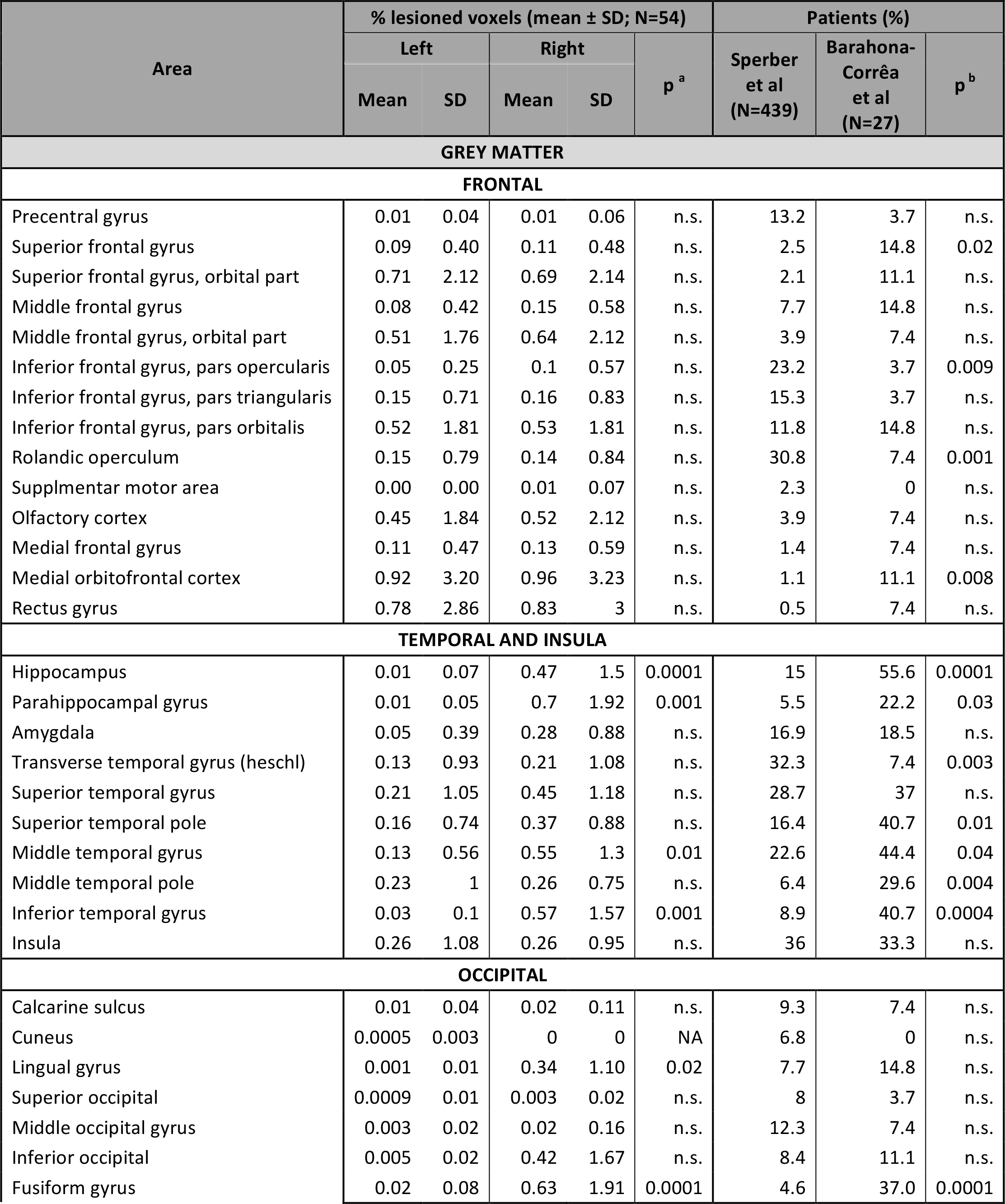
Comparison of Right Hemisphere Lesion with Left Hemisphere Lesions and with a Control Sample of Stroke Lesions

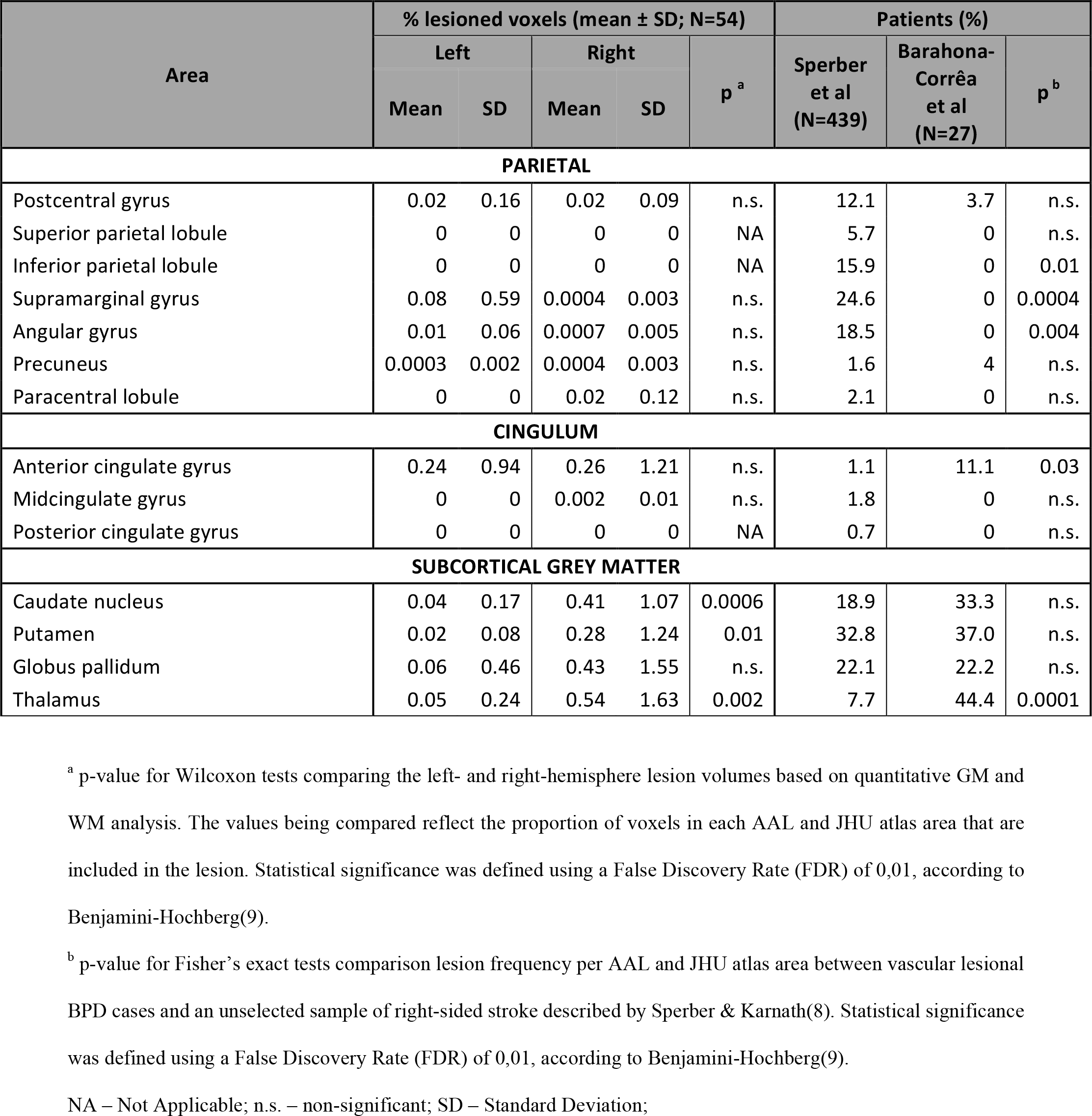

To further confirm the relevance of specific brain regions, we then compared lesion distribution among specific sub-samples of our database with that of patients with lesions of similar etiology, but not selected for any particular symptom. Compared to the distributions of brain tumors described by Ostrom et al(7), tumors associated with lesional mania were more frequently located in the temporal lobes, brain stem, and cerebrum, which includes the thalamus and basal ganglia (Fig. 2D). Among lesional mania patients with traced right-sided vascular lesions, a more detailed comparison with MRI scans from 439 right-hemisphere stroke patients described by Sperber & Karnath (8) showed that lesional mania patients were more frequently affected in the superior frontal gyrus, medial orbitofrontal cortex, hippocampus and parahippocampal gyrus, superior and middle temporal poles, middle and inferior temporal gyrus, fusiform gyrus, anterior cingulate gyrus, and thalamus (table 2). In contrast, the inferior frontal gyrus, rolandic operculum, transverse temporal gyrus, and several parietal regions were less frequently involved in lesional mania cases.

Finally, we performed exploratory sensitivity analyses, restricted only to cases of lesional mania secondary to vascular lesions, with no personal or family history of neuropsychiatric disorder, and with available CT or MRI images. These analyses confirmed predominance of right-sided vs. left-sided lesions in patients with lesional mania, overall (78.9% vs. 34.2%; p<0.01, McNemar’s test) and in the putamen, thalamus and hippocampus. In the two latter areas, as well as the middle temporal pole and fusiform gyrus, an excess of lesions was found in lesional mania patients, when compared with right-hemisphere stroke control patients (Table S5).

## Discussion

The main aim of this work was to test the hypothesis that lesional mania is more often associated with right-hemisphere than left-sided lesions. Towards this aim, we performed a systematic literature review of case-reports and case-series, followed by pooled analysis of individual patient and lesion data from eligible cases. We retrieved 207 cases of lesional mania, most of which with right-hemisphere focal brain lesions (84.3%), and only 34.5% with left-sided lesions. This difference was statistically significant, and was conserved across different disease subtypes as well as in analyses restricted to patients without personal or family history of neuropsychiatric disorder. To our knowledge, this is the first systematic review and pooled analysis of published cases of lesional mania, offering the most comprehensive demonstration of a long-held, albeit empirically unconfirmed, axiom of textbook neuropsychiatry, i.e., that lesional mania is preferentially associated with right-hemisphere, rather than left-hemisphere, brain lesions.

Our main result confirms previous works based on smaller samples(12, 13). Robinson and colleagues found a predominance of right-sided brain lesions in seventeen lesional mania cases when compared to thirty-one patients with post-stroke depression, who had predominantly left-sided lesions. Most lesional mania patients had lesions involving the right orbitofrontal and basotemporal cortex, caudate and thalamus, while patients with post-stroke depression had more widely distributed lesions predominantly involving the head of the left caudate and the left insular and basotemporal cortex. Moreover, there was no overlap between right-sided lesions associated with mania and right-sided lesions associated with depression. Starkstein and colleagues reported similar findings in an independent cohort of eight lesional mania patients and further complemented the anatomical analysis with results from ^18^fluorodeoxyglucose positron emission tomography performed in three patients, which showed a lower ^18^fluorodeoxyglucose uptake in several right limbic regions including lateral basotemporal and superior frontal areas.

Additionally, our main finding converges with recent voxel-based meta-analyses of GM and WM abnormalities in primary BPD, which point to significant laterality effects. For instance, in a meta-analysis of eight voxel-based morphometry studies that compared BPD patients and healthy controls, Selvaraj and colleagues found a right-sided contiguous cluster of GM reduction in BPD patients compared to controls, encompassing the insula, middle and superior temporal gyrus, temporal pole, pars opercularis and pars triangularis, inferior frontal gyrus, and claustrum(14). More recently, two other voxel-based meta-analyses by Wise and colleagues described clusters of lower GM volume in the right middle and inferior temporal gyri and right middle occipital gyrus in patients with BPD, as well as decreased fractional anisotropy in the right anterior superior longitudinal fasciculus(15, 16). Functional MRI studies in adults with BPD have also shown predominantly right-sided decreased signal in the ventrolateral prefrontal emotional arousal network and associated limbic structures such as the parahippocampus and lingual gyrus, during tasks that require processing of emotional stimuli or adaptive response-inhibition(17, 18). Moreover, in another study, manic symptoms significantly correlated with activation of the right amygdala, right inferior frontal gyrus and right putamen(18). Resting-state fMRI studies have likewise shown decreased functional connectivity strength in right supramarginal gyrus and angular gyrus, right superior frontal gyrus, and right superior parietal gyrus(19). Finally, EEG records of hypomanic BPD patients have shown higher relative left-frontal alpha-power compared to healthy subjects(20), adding to the evidence for a right-left imbalance in brain function for BPD patients.

Our analyses further suggest that right-hemisphere predominance of lesions in lesional mania follows a non-random anatomical distribution, resulting mostly from lesions of specific areas, namely the hippocampus, parahippocampal gyrus, middle and inferior temporal gyri with adjoining WM, lingual and fusiform gyri, caudate nucleus, putamen and thalamus, and both limbs of the internal capsule (Table 2; Table S4). Importantly, we also found significant differences in lesion distribution when comparing lesion mania subsamples with large samples of patients with brain lesions of a similar nature, but not selected for particular behavioral outcomes. Specifically, in comparisons with patients with right-hemisphere stroke(8), right-sided vascular lesions associated with lesional mania affected more frequently all of these areas, with the exception of the lingual gyrus, caudate nucleus, putamen and WM areas. This suggests that right-sided predominance of lesions is not merely attributable to a laterality bias in stroke incidence in the general population. Moreover, while the limited quality of lesion topography depiction in most case-reports advises caution in interpreting the over-representation of these areas in lesional mania, it is nevertheless remarkable that all these areas have been repeatedly highlighted by structural or functional neuroimaging studies in primary BPD, often with right-sided lateralized findings(21–23).

In addition to the thalamus, temporal lesions, in particular those affecting the right hippocampus and parahippocampal gyrus, were the most consistently over-represented in lesional mania, both in left-right comparisons and in comparison with tumor and vascular controls. In fact, there is reasonable consensus regarding hippocampal abnormalities in primary BPD, but it remains unclear if they reflect treatment effects or disease progression, rather than neural vulnerability for the disorder(21). Our results support the latter. Occipital cortical areas, fusiform and lingual gyri in particular, were also over-represented in right-sided lesions, with the fusiform gyrus also more frequently involved in lesional mania when compared to stroke controls. While occipital areas are seldom mentioned in BPD literature, at least two independent fMRI studies found fusiform gyrus hypoactivation during emotional face processing in primary BPD(24, 25).

Several areas, namely the right temporal pole and several frontal lobe areas, were identified in comparisons with the control stroke sample, but not the left-right comparison (table 2). The temporal pole is part of the right-sided contiguous cluster of GM volume reduction in primary BPD patients identified by Selvaraj et al (2012), and there was a recent report of reduced diffusivity entropy in the temporal poles among these patients(26). Regarding frontal areas, lesions in Brodmann area 11 and the anterior cingulate are of particular interest, since functional abnormalities have been consistently reported for these areas in primary BPD(17, 22, 23) and the anterior cingulum was part of the cluster of areas where Wise and colleagues found significantly reduced GM volumes(15). It is noteworthy that the OFC and anterior cingulate did not stand out in the left/right comparison, since findings of morphometric GM studies in primary BPD consistently report lower GM volumes in these areas, but no laterality effects. Absent lateralization regarding frontal areas could in fact reflect their critical role, with possibly similar functional effects resulting either from a lesion in a particular hemisphere, or from disturbed frontal interhemispheric connectivity secondary to a lesion in the contralateral hemisphere (27). Finally, as has been reported in other lesion studies (28), several areas were less frequently affected in vascular lesional mania compared to the vascular control group (8), possibly reflecting differing vascularization patterns for these areas and those that are over-represented in lesional mania.

The wide distribution of GM areas highlighted by our analyses argues for a potential circuit-based impact of the several different lesions associated with secondary mania. In fact, some of these areas, namely the superior frontal gyrus, including its orbital part, anterior cingulate, hippocampus/parahippocampal gyrus, and inferior temporal gyrus, partly overlap with the Default Mode Network (DMN)(29). Gray-matter volume reduction has been found among primary BPD patients in several components of the DMN, namely the prefrontal cortex(23, 30), cingulate cortex(22, 31), temporal gyri(31) and hippocampus(22). Consistently, in primary BPD, functional connectivity studies show reduced coherence and connectivity strength in several DMN nodes(19), and there is evidence of a left-predominant asymmetry of the DMN(32), further reinforcing the validity of a preferentially right-sided distribution of brain lesions in lesional mania. In any case, the wide distribution of areas associated with lesional mania merits further analyses, possibly using novel approaches for network localization of symptoms from focal brain lesions(33), that have already been used for analysis of other lesional neuropsychiatric syndromes(34).

Our findings, while novel and informative, should be interpreted considering the limitations of the study design. First and foremost, our analyses were restricted to author descriptions of lesions and/or a limited number of slices from each scan, limiting the accuracy of lesion mapping and the validity of topographical analysis. Moreover, the choice of published scan slices by authors could be potentially biased by expectations regarding lesion location. Irrespective of these limitations, it is important to note that right-hemisphere predominance was restricted to many of the same areas that were identified in the comparison with the stroke control group, cross-validating these findings and suggesting that, for these structures, a specific interaction does exist between laterality and mood regulation. Importantly, authors and reviewers may also have been biased towards publication of cases confirming the conventional view that secondary BPD is associated with right hemisphere lesions. Nevertheless, the contrary could also be true, considering the tendency to publish rare associations. In any event, among published reports of lesional mania, the number of right-sided lesions has been consistently higher than left-sided lesions since before the 1970’s, arguing against the existence of such a bias (see figure S1 for details). Another possible bias may result from under-diagnosis of milder hypomanic syndromes in aphasic patients with left-hemisphere lesions. However, right-hemisphere predominance in our analysis was not restricted to brain areas involved in language production, which does not support this hypothesis.

In conclusion, our study provides the most solid demonstration to date of the association between right-hemisphere brain lesions and the development of secondary mania, while suggesting the first systematic mapping of lesion topography in lesional mania. In that respect, we found that in lesional mania, specific brain areas, distributed across multiple brain regions and circuits, are most frequently affected. We expect these findings will contribute to a more thorough understanding of the role of these brain areas in mood regulation and their importance in the context of bipolar disorders, specifically with regards to lateralization in the control of such functions and in the development of these disorders.

## DECLARATION OF INTEREST

None of the authors have declared potential conflicts of interest involving this work, including relevant financial activities outside the submitted work and any other relationships or activities that readers could perceive to have influenced, or that give the appearance of potentially influencing what is written.

## FUNDING

GC was supported by Fundação para a Ciência e Tecnologia (FCT) through a PhD Scholarship (SFRH/BD/130210/2017). AJO-M was supported by FCT through a Junior Research and Career Development Award from the Harvard Medical Portugal Program (HMSP/ICJ/0020/2011). FCT did not have a role in the design and conduct of the study, in the collection, management, analysis, and interpretation of the data, in the preparation, review, or approval of the manuscript, nor in the decision to submit the manuscript for publication.

## AUTHOR CONTRIBUTION

JBB-C, GC, RMC and AJO-M conceived and designed the work; GC, RR, AV, VCS, CS, H-OK and SS acquired the data; JBB-C, GC, CS, H-OK, SS and AJO-M, analyzed and interpreted data; JBB-C, GC and AJO-M drafted the work; RMC, RR, AV, VCS, CS, H-OK and SS revised the manuscript critically for important intellectual content; all authors approved the final version to be published and agree to be accountable for all aspects of the work in ensuring that questions related to the accuracy or integrity of any part of the work are appropriately investigated and resolved.

